# Time-Dependent Material Properties and Composition of the Nonhuman Primate Uterine Layers Through Gestation

**DOI:** 10.1101/2024.11.17.624020

**Authors:** Daniella M. Fodera, Echo Z. Xu, Camilo A. Duarte-Cordon, Michelle Wyss, Shuyang Fang, Xiaowei Chen, Michelle L. Oyen, Joy Y. Vink, Ivan Rosado-Mendez, Helen Feltovich, Timothy Hall, Kristin M. Myers

**Affiliations:** Department of Biomedical Engineering, Columbia University, New York, NY, USA; Department of Mechanical Engineering, Columbia University, New York, NY, USA; Department of Biomedical Engineering, Virginia Tech, Blacksburg, VA, USA; Department of Pathology and Cell Biology, Columbia University Irving Medical Center, New York, NY, USA; Department of Biomedical Engineering, Wayne State University, Detroit, MI USA; Department of Obstetrics & Gynecology, John A. Burns School of Medicine, University of Hawai’i at Maā noa, Honolulu, HI, USA; Department of Medical Physics, University of Wisconsin-Madison, Madison, WI, USA; Department of Radiology, University of Wisconsin-Madison, Madison, WI, USA; Department of Obstetrics & Gynecology, North Memorial Health System, Robbinsdale, MN, USA

**Keywords:** uterus, pregnancy, rhesus macaque, microindentation, poro-viscoelasticity, reproductive biomechanics

## Abstract

The uterus is central to the establishment, maintenance, and delivery of a healthy pregnancy. Biomechanics is an important contributor to pregnancy success, and alterations to normal uterine biomechanical functions can contribute to an array of obstetric pathologies. Few studies have characterized the passive mechanical properties of the gravid human uterus, and ethical limitations have largely prevented the investigation of mid-gestation periods. To address this key knowledge gap, this study seeks to characterize the structural, compositional, and time-dependent micro-mechanical properties of the nonhuman primate (NHP) uterine layers in nonpregnancy and at three time-points in pregnancy: early 2^nd^, early 3^rd^, and late 3^rd^ trimesters. Distinct material and compositional properties were noted across the different tissue layers, with the nonpregnant endometrium and pregnant decidua being the least stiff, most viscous, least diffusible, and most hydrated layers of the NHP uterus. Pregnancy induced notable compositional and structural changes in the decidua and myometrium but had no effect on their micro-mechanical properties. Further comparison to published human data revealed marked similarities across species, with minor differences noted for the perimetrium and nonpregnant endometrium. This work provides insights into the material properties of the NHP uterus and demonstrates the validity of NHPs as a model for studying certain aspects of human uterine biomechanics.

## Introduction

The biomechanical functions of the female reproductive system critically underpin the dynamic physiologic processes of pregnancy^1,2^. The uterus, in particular, undergoes dramatic growth and remodeling in pregnancy to enable fetal growth and development^3–6^. Biomechanical defects to this organ, at the cell and tissue length scales, are thought to cause an array of obstetric disorders, including, but not limited to, preterm birth, intrauterine growth restriction, and uterine rupture, which in turn contribute to the high incidence of maternal and fetal morbidity and mortality in the United States^1,7–11^. To elucidate the role of biomechanics in the pathogenesis of obstetric conditions, a baseline knowledge of normal uterine structure, composition, and mechanics throughout the course of pregnancy is first needed.

Anatomically, the human uterus is an inverted, pear-shaped organ with a single uterine cavity^1,3,12^. The uterine wall is composed of three structurally and functionally distinct tissue layers: (i) the nonpregnant endometrium and pregnant decidua, (ii) the myometrium, and (iii) the perimetrium (i.e., serosa)^1,3,12^. The nonpregnant endometrium and pregnant decidua together represent the innermost uterine layer consisting of luminal and glandular epithelial cells, stromal cells, and spiral arteries embedded in a collagen-dense extracellular matrix (ECM)^3,13^. In nonpregnancy, the endometrium undergoes cyclic cellular and molecular changes throughout the menstrual cycle in response to hormonal fluctuations^13^. The decidua, the pregnant counterpart of the endometrium, forms the basis of the maternal-fetal interface, providing critical nutritional support and immunological protection for the developing fetus^3^. The myometrium, the middle and thickest layer of the uterus, is primarily composed of smooth muscle fascicles interwoven with collagen and elastin fibers and pocketed with blood vessels^3^. Throughout pregnancy, the myometrium must undergo passive growth and stretch to accommodate the growing size of the fetus through smooth muscle cell hyperplasia and hypertrophy^3^. In addition to passive mechanical functions, the myometrium exhibits active contractile behavior to enable sperm motility and menstrual blood egress in nonpregnancy and forceful uterine contractions during labor^3^. Exterior to the myometrium and adjacent to the abdominal cavity is the perimetrium (i.e., serosa), a thin collagen-dense tissue layer that acts as a smooth, lubricated barrier for the uterus^3,12^.

The pregnant human uterus is a protected environment, and ethical considerations limit deep structure-function investigations to two distinct physiologic stages: nonpregnancy and late 3^rd^ trimester^1^. To overcome this barrier, animal models have been previously used to interrogate mid-gestational changes to maternal and fetal physiology, however, gross reproductive anatomy and pregnancy characteristics differ dramatically across mammalian species^14–16^. The anatomic and physiologic similarities of Rhesus macaques (*Macaca mulatta*) and humans are well established^17,18^. With regards to the uterus, both species have three distinct uterine layers surrounding a single uterine cavity, undergo menstruation in nonpregnancy, and most often carry singleton pregnancies to term (Fig. 1)^17,18^. Humans and Rhesus macaques notably differ in total gestational length (270 vs 160 days), depth of embryo implantation, degree of decidualization during the menstrual cycle, number of placental discs, overall lifespan (70 vs 30 yrs), and method of locomotion (bipedal vs quadrupedal)^3,17,18^.

**Figure 1.**
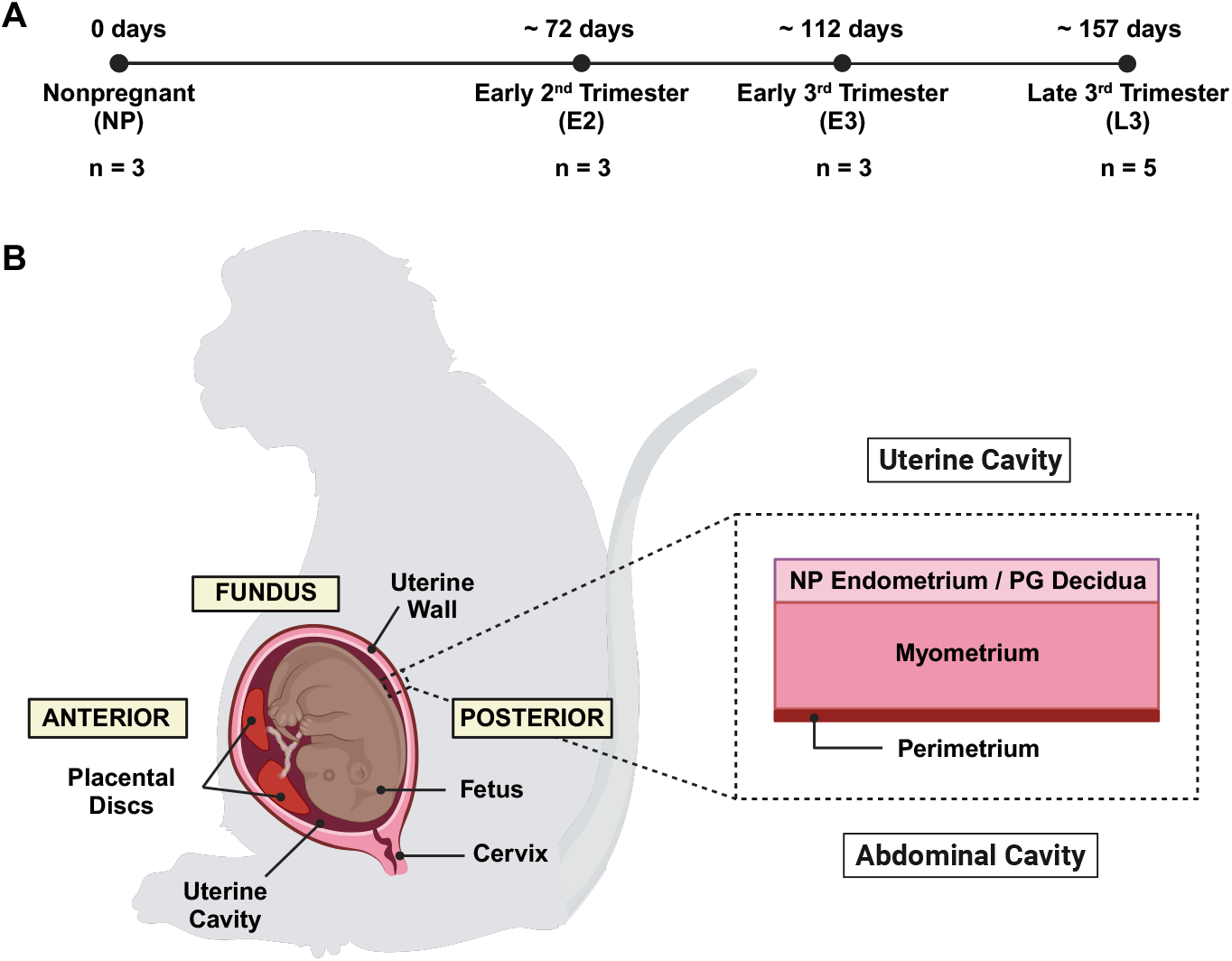
Pregnant Rhesus Macaque Anatomy. **(A)** Timeline of Rhesus macaque pregnancy. **(B)** Representative illustration of pregnant Rhesus macaque anatomy in the sagittal plane. Relevant reproductive structures and anatomic regions (anterior, fundus, and posterior) are labeled, including a detailed schematic of the uterine wall containing all three uterine layers (NP endometrium/ PG decidua, myometrium, perimetrium).

Previous work has characterized the passive material properties of the human uterus at multiple length scales, yet no studies to date have evaluated the mechanics of the nonhuman primate (NHP) uterus^19–27^. On the micrometer length scale, microindentation has been previously employed by our group to measure the time-dependent material properties of all three uterine layers for humans in nonpregnancy and late 3^rd^ trimester^20^. Significant variations in all material properties were noted across tissue layers, with the endometrium and decidua being the least stiff, most viscous, and least permeable^20^. In human pregnancy, the decidua exhibited increases in stiffness, viscoelastic ratio, and diffusivity, while no changes were observed for the myometrium or perimetrium^20^. Further, a study by Abbas et al. (2019) measured the stiffness of nonpregnant endometrium and first-trimester decidua tissues with atomic force microscopy and noted no change in stiffness between these tissue types at the micro-scale^19^. For larger testing regimes on the millimeter to centimeter length scale, studies have exclusively characterized the myometrium in nonpregnancy and late pregnancy using tension, compression, indentation, and shear^21–27^. Overall, the human myometrium exhibits nonlinearity, anisotropy, and tension-compression asymmetry, with nonpregnant tissue exhibiting increased stiffness and decreased extensibility compared to pregnant tissue^21–27^.

It is presently unknown how the material and structural properties of the human uterus change in a healthy pregnancy between the first and third trimesters. Therefore, we seek to utilize a nonhuman primate (NHP) model to characterize mid-gestational changes to the mechanical and structural properties of the uterus, distinguishing across all three tissue layers (Fig. 1). Specifically, this study will investigate nonpregnant (NP) and pregnant (PG) states in early 2^nd^ (E2), early 3^rd^ (E3), and late 3^rd^ (L3) trimesters. We expect that mechanical and structural changes observed in the NHP model will mimic trends noted previously for humans and enable a more complete biomechanical understanding of pregnancy.

## Results

### Structure and Composition of NHP Uterine Layers

The structure and composition of all three uterine tissue layers (i.e., NP endometrium / PG decidua, myometrium, and perimetrium) were evaluated from NHP subjects (i.e., Rhesus macaques) in nonpregnancy (N = 3) and pregnancy at E2 (N = 3), E3 (N = 3), and L3 (N = 5) trimesters (Fig. 1). All tissues were reviewed by a board-certified pathologist; menstrual cycle stage and pathological findings were noted in Table S1. The uterine tissue layers of the NHP exhibited distinct structure and composition of ECM and cellular components (Fig. 2A). In nonpregnancy, the endometrium was primarily composed of pseudo-stratified epithelial glands and densely-packed stromal cells with a small portion of immune cells (Fig. 2A). Blood vessels comprised less than 10% of the overall endometrial tissue area and were concentrated in the basalis layer immediately adjacent to the myometrium (Fig. 2E). Collagen was diffusely present in the stromal spaces of the endometrium and tightly surrounded the endometrial glands to act as a basement membrane (Fig. 2A). Compared to the superficial functionalis layer of the endometrium, increased deposition of collagen was found in the basalis layer.

**Figure 2.**
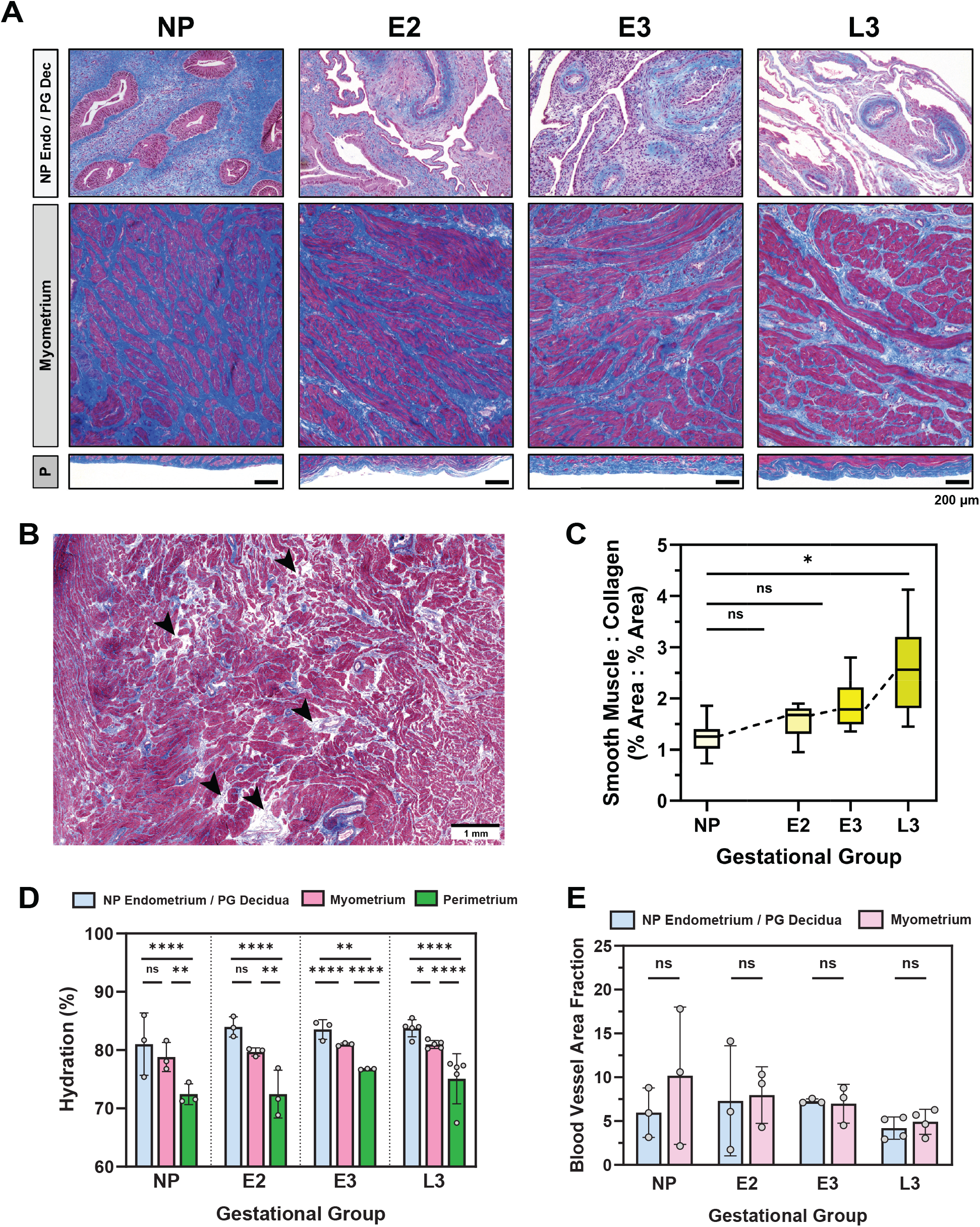
**(A)** Histology of NHP uterine layers [NP endometrium (Endo) / PG decidua (Dec), myometrium, and perimetrium (P)] across gestation. Tissues are stained with Masson’s Trichrome (blue = collagen; red = smooth muscle, cytoplasm; black = nuclei). Note that the relative lengths of the tissue layer figure panels do not reflect actual layer proportions. **(B)** Representative image of focal edema in the L3 myometrium. **(C)** Ratio of smooth muscle to collagen content in the myometrium across gestation. All animal subjects, evaluated at a single anatomic region, are represented in these data. **(D)** Tissue hydration of all uterine layers across gestation sampled at the posterior region. All animal subjects (*n* = 3−5) in this study are represented in these data. **(E)** Blood vessel area fraction in the NP endometrium, PG decidua, and myometrium for each gestational group. All animals, except one L3 subject, are represented in these data. **22**

In pregnancy, the endometrium dramatically remodels into the decidua. All decidua tissue taken from NHPs in this study can be classified as decidua parietalis, distant from the sites of placentation. Overall, the epithelial glands appeared flattened, and decidualized stromal cells adopted a polygonal shape (Fig. 2A). Blood vessels continued to be present in the PG decidua, both superficially and deep, but displayed no notable changes in size and concentration relative to nonpregnancy (Fig. 2E). Collagen was diffusely present throughout the decidua tissue and was concentrated around blood vessels (Fig. 2A). No notable changes to the structure and composition of the decidua were observed across E2, E3, and L3 gestational groups (Fig. 2A).

The myometrium, the middle and thickest layer of the uterus, exhibited longitudinal alignment of smooth muscle fibers surrounded by thick bands of collagen for all NHP subjects (Fig. 2A). Blood vessels represented, on average, 10% or less of the overall myometrial tissue area; the largest blood vessels appeared centered in the middle third of the uterine wall (Fig. 2E). In pregnancy, the smooth muscle cells of the myometrium underwent hypertrophy, exhibiting an increase in cell volume. The relative proportion of smooth muscle to collagen content increased in late third trimester relative to nonpregnancy as determined through semi-quantitative image analysis (Fig. 2C). No change in the distribution and size of blood vessels was noted for the myometrium in pregnancy (Fig. 2E). Interestingly, a unique phenomenon of focal edema was observed for all L3 pregnant tissues evaluated: increased interstitial spacing between the collagen and smooth muscle cells (Fig. 2B). This histological feature was not observed for either E2 or E3 groups and given its localized nature and consistency of appearance for all L3 tissues, it is unlikely to be the product of a histological artifact.

Lastly, the perimetrium, also known as the serosa, appeared as a thin, smooth band of collagen adjacent to the myometrium (Fig. 2A). In a subset of samples, thicker regions of collagen indicative of fibrosis and small amounts of vasculature were visible in the perimetrium (Table S1). No overt changes to this tissue layer as a result of pregnancy were noted (Fig. 2A).

In addition to histological analysis, the hydration of each uterine layer was quantified by means of lyophilization. Tissue hydration was determined to be distinct across uterine tissue layers (endometrium/decidua: 83.2 ± 2.7%; myometrium: 80.2 ± 1.4%; perimetrium: 74.3 ± 3.5%), with the perimetrium being the least hydrated tissue layer of the uterus (Fig. 2D). No change in hydration was observed across gestation for any uterine tissue layer (Fig. 2D).

### Material Properties of NHP Uterine Layers

Spherical microindentation (*R* = 50 *µ*m) was employed in this study to measure the time-dependent material properties of NHP uterine layers, namely the NP endometrium, PG decidua, myometrium, and perimetrium, across gestation (Fig. 3A). Tissues were taken from three anatomic regions (i.e., anterior, fundus, and posterior) from the same NP, E2, E3, and L3 animal subjects described previously away from visible sites of pathology (Fig. 1). Approximately 100 indentation points were measured for each tissue sample, representing more than 12,000 individual indentation measurements in all. To describe the uterus’ intrinsic viscoelasticity (rearrangement of the solid matrix) and poroelasticity (fluid flow migration), phenomena known to be exhibited by soft biological tissues^28–30^, an established poroelastic-viscoelastic (PVE) constitutive model^31^ was employed to determine the following material parameters: instantaneous elastic modulus (*E*_0_), equilibrium elastic modulus (*E*_∞_), poroelastic modulus (*E*_*PE*_), viscoelastic ratio (*E*_∞_*/E*_0_), intrinsic permeability (*k*), and diffusivity (*D*). Representative force versus indentation depth and force versus time curves are shown in Figs. 3B and C.

**Figure 3.**
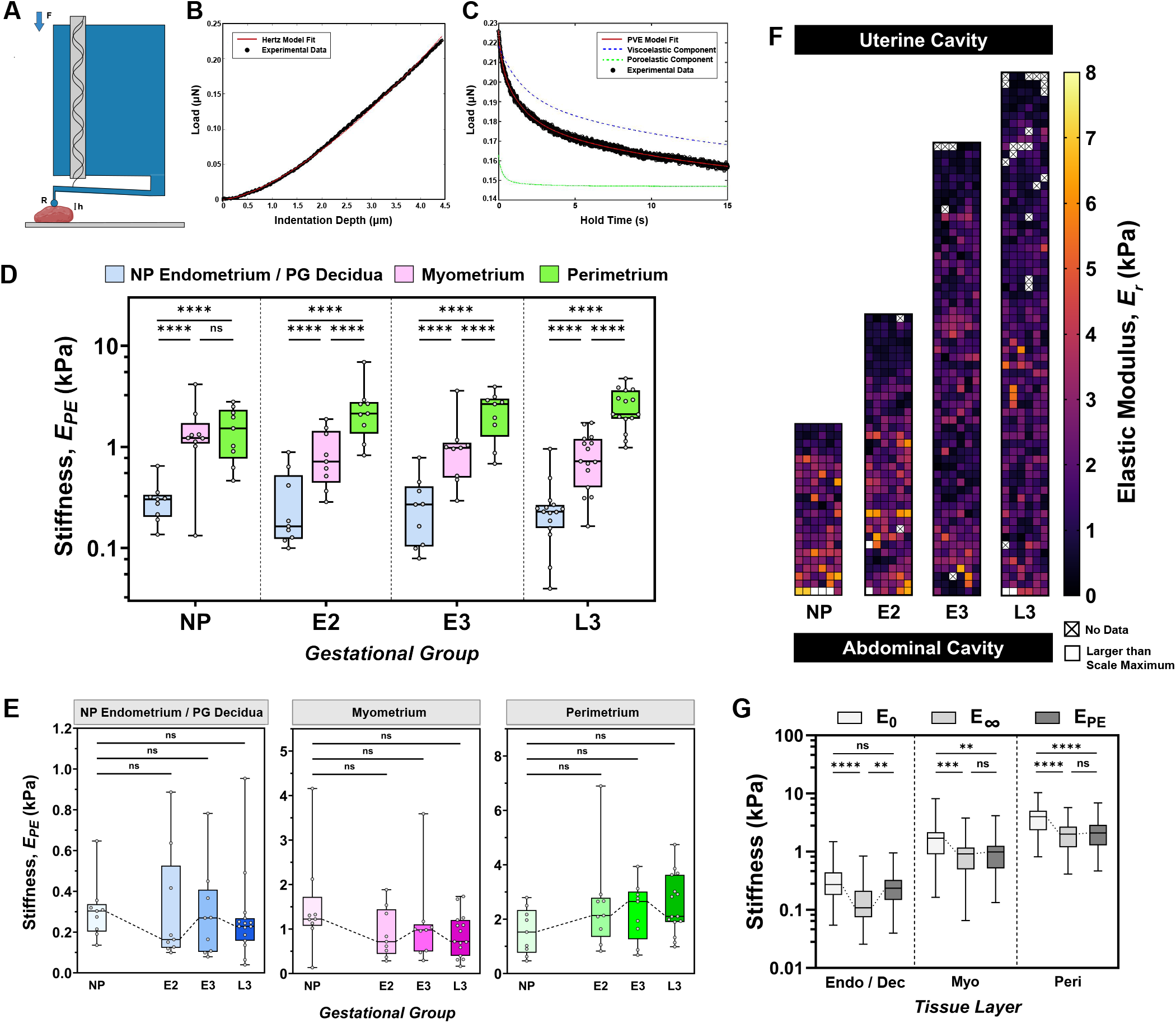
**(A)** Schematic of the Piuma nanoindenter (Adapted from Optics11 Life). A spherical probe with radius (*R*) is attached at the end of a cantilever and indented into the sample at a fixed depth (*h*), recording load (*F*) over time. **(B)** Representative load vs indentation data fitted with the Hertzian contact model. **(C)** Representative load vs time data fitted with the PVE model. **(D-E)** Elastic modulus (*E*_*PE*_) of the NHP uterus **(D)** across tissue layers and **(E)** across gestation. Each point represents the median value of all indentation points measured for a single sample. All animal subjects (*n* = 3 − 5) in this study sampled at three anatomic regions (anterior, fundus, posterior) are represented in these data. **(F)** Spatial variation in local elastic modulus/stiffness (*E*_*r*_) values across the uterine walls of NP, E2, E3, and L3 subjects. Measurements were taken at every 200 µm across the length of the tissue. Points removed due to exclusion criteria are represented by [X]. White squares indicate data points outside the bounds of the y-axis. **(G)** Comparison of viscoelastic (*E*_0_, *E*_∞_) and poroelastic (*E*_*PE*_) elastic modulus parameters for each tissue layer with data being pooled from all gestational groups and anatomic regions.

Surprisingly, no changes in any of the material parameters were observed across gestation for the NP endometrium, PG decidua, myometrium, and perimetrium tissue layers (Fig. 3E, Fig. 4B,D,F). The greatest differences in material properties were found across distinct uterine layers for each gestational group evaluated (Fig. 3D). All elastic modulus parameters (*E*_0_, *E*_∞_, and *E*_*PE*_), which are measures of tissue stiffness (resistance to deformation), ranged from 10^1^ to 10^4^ Pa, were highly correlated with one another, and exhibited identical trends across tissue layers and gestational groups (Fig. S4). Overall, tissue stiffness increased from the intra-uterine cavity to the outer abdominal cavity, with the endometrium and decidua being the least stiff and the perimetrium being the most stiff (Fig. 3D). The perimetrium was stiffer than the NP endometrium and PG decidua layers for all gestational groups evaluated; only for PG time points was the perimetrium stiffer than the myometrium (Fig. 3D). Spatial variations in tissue stiffness (*E*_*r*_) were assessed across the entire uterine wall thickness (Fig. 3F). Notably, a stiffness gradient was observed at the interfaces between the NP endometrium and myometrium, as well as between the PG decidua and myometrium (Fig. 3F). Across the individual elastic modulus parameters measured, instantaneous elastic modulus (*E*_0_), as expected, was greater than the equilibrium elastic modulus (*E*_∞_) for all samples (Fig. 3G). Between *E*_∞_ and *E*_*PE*_, no difference was observed for the myometrium and perimetrium tissue layers, but there was a minute but systemic increase in *E*_*PE*_ relative to *E*_∞_ for the NP endometrium and PG decidua layers (Fig. 3G).

**Figure 4.**
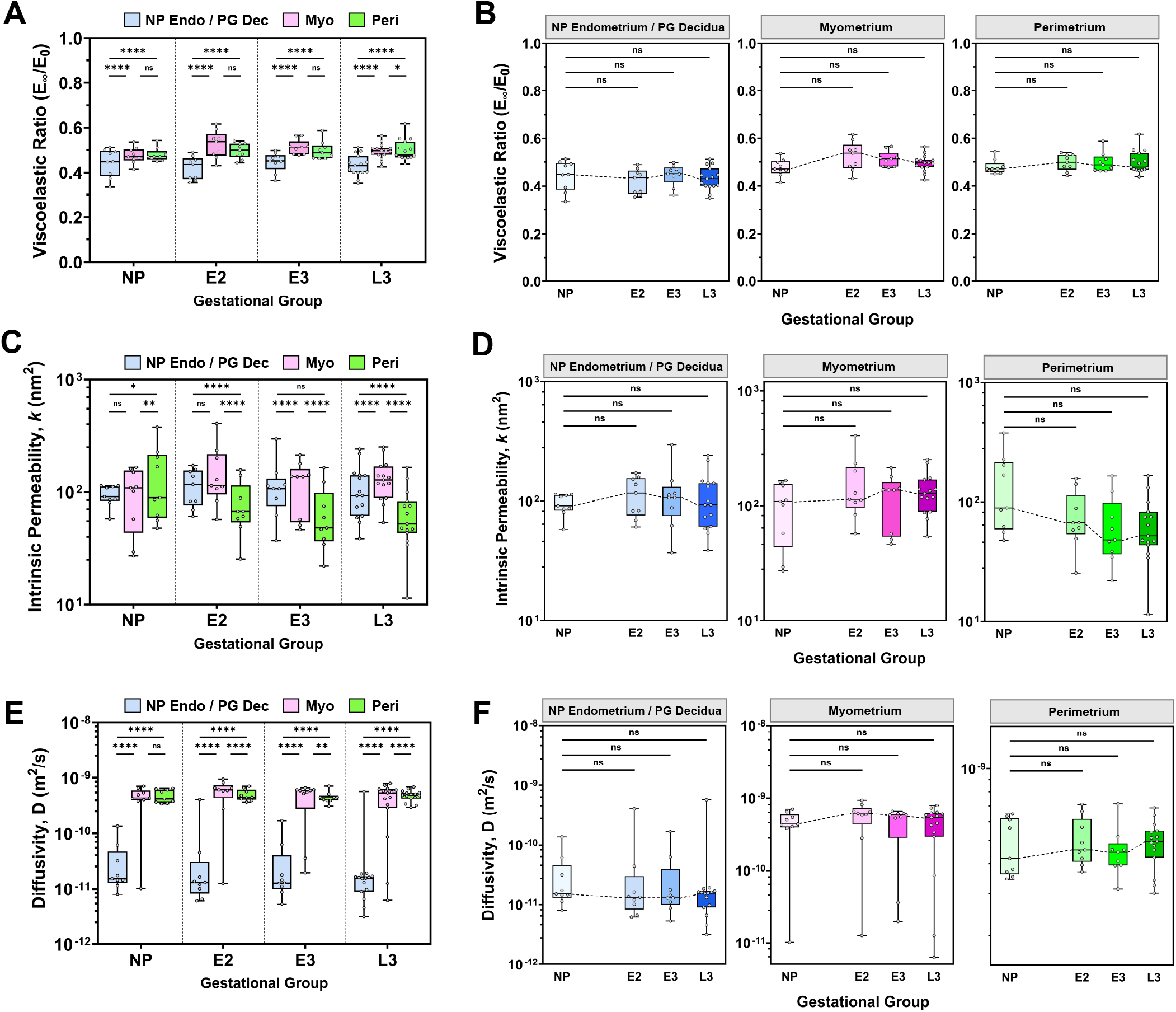
Viscoelastic ratio **(A, B)**, intrinsic permeability **(C, D)** and diffusivity **(E, F)** of the NHP uterus **(A, C, E)** across tissue layers [Endo = Endometrium, Dec = Decidua, Myo = Myometrium, Peri = Perimetrium] and **(B, D, F)** across gestation. Each point represents the median value of all indentation points measured for a single sample. All animal subjects (*n* = 3−5) in this study sampled at three anatomic regions (anterior, fundus, posterior) are represented in these data.

Median values of viscoelastic ratio (*E*_∞_*/E*_0_) ranged between 0.3 and 0.6 for all samples evaluated, indicating that the uterus possesses both solid-like and fluid-like material behavior (Fig. 4A,B). The NP endometrium and PG decidua layers were determined to be slightly more viscous (0.43 ± 0.05) than the myometrium (0.50 ± 0.04) and perimetrium (0.49 ± 0.04) layers for all gestational groups (Fig. 4A). No statistically significant difference in viscoelastic ratio was observed between the myometrium and perimetrium layers except in the L3 group (Fig. 4A). Intrinsic permeability (*k*) is an innate property of a porous medium (e.g., biological tissue) that describes a material’s resistance to fluid flow as a product of its pore geometry. Values of intrinsic uterine permeability ranged between 10^1^ to 10^3^ *nm*^2^ for all tissue layers (Fig. 4C,D). For all gestational groups, the permeability of the perimetrium (87 ± 68 *nm*^2^) was slightly less than the endometrium/decidua (110 ± 53 *nm*^2^) and myometrium (131 ± 70 *nm*^2^) layers (Fig. 4C). Slight variations in permeability values between the endometrium/decidua and myometrium layers occurred only for E3 and L3 groups (Fig. 4C). At this length scale of material testing, average pore size (*ξ*), whereby 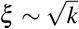, was determined to be in the range of 4 to 14 nm. Lastly, diffusivity (*D*), also known as the diffusion coefficient, is a measure that describes the flow of fluid through a porous medium over time. For the uterus, diffusivity is significantly decreased in the NP endometrium and PG decidua layers, by more than an order of magnitude (0.45 ± 1.07 *x* 10^−10^ *m*^2^*/s*), when compared to the myometrium (4.79 ± 2.43 *x* 10^−10^ *m*^2^*/s*) and perimetrium (4.82 ± 1.13 *x* 10^−10^ *m*^2^*/s*) layers (Fig. 4).

The effect of tissue and subject characteristics on the material properties of the uterus was also investigated in this study. Across the three anatomic regions evaluated (i.e., anterior, posterior, and fundus), regional variations in all material properties occurred on an individual animal basis for each of the tissue layers and gestational groups (Fig. S2). However, it remains unclear whether these differences represent consistent systemic trends across all NHP subjects due to small sample size considerations. In addition, no material properties reported in this study correlated linearly with animal age and gravidity, defined as the total number of previous pregnancies (Fig. S3). To note, age and gravidity were considered together in the linear regression model since, in this cohort of NHPs studied, there was an increasing linear correlation between animal age and gravidity (Fig. S1).

Further investigation into the inter-correlation of material properties revealed a unique scattering of data between tissue stiffness and permeability, which was characteristic of each distinct uterine tissue layer (Fig. 5B). Notably, the perimetrium exhibits one primary cluster of data with a negative linear relationship between stiffness and permeability on a log-log scale, while the myometrium displays two distinct, linearly aligned data clusters (Fig. 5A). No relationship between permeability and stiffness exists for the endometrium and decidua tissue layers (Fig. 5, S4).

**Figure 5.**
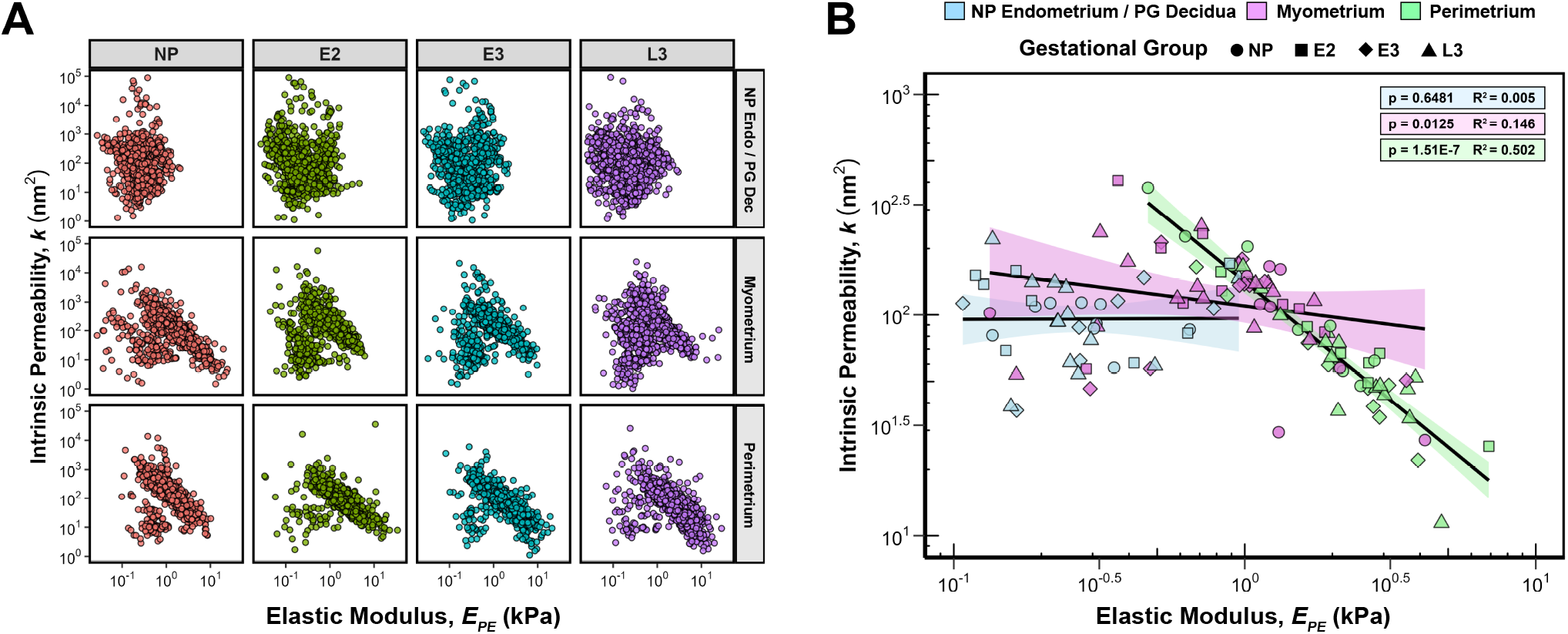
Correlation of elastic modulus (*E*_*PE*_) and intrinsic permeability (*k*) parameters for **(A)** all data points, separated by tissue layer and gestational group, and **(B)** median values for each sample, separated by tissue layer.All animal subjects (*n* = 3−5) in this study sampled at three anatomic regions (anterior, fundus, posterior) are represented in these data. Linear regression analysis was performed for each tissue layer with data being pooled across all gestational groups. *R*^2^ and *p* values are noted; shaded regions indicate the 95% confidence interval.

### Comparative Analysis of Human and Rhesus Macaque Uterine Layer Material Properties

Microindentation data generated by this study on NHP uterine layers was directly compared to published data for the human uterus that employed similar methodologies^19,20^. Nonpregnant and late third-trimester (PG-CS) data on all uterine layers were sourced from Fodera et al. (2024), while first-trimester decidua data were taken from Abbas et al. (2019). Notable similarities and differences in the time-dependent material properties of the human and NHP uterus were found in nonpregnancy and late third trimester (Fig. 6). Comparing between humans and NHPs, no significant differences in the values of viscoelastic ratio, permeability, and diffusivity were found for all three uterine tissue layers for NP and L3 PG time points (Fig. 6). Interestingly, the relative changes between NP and PG groups were notably different for the NP endometrium and PG decidua layers. In humans, there was a statistically significant increase in the viscoelastic ratio, permeability, and diffusivity parameters for the PG decidua relative to the NP endometrium^20^. Such trends did not exist for NHPs.

**Figure 6.**
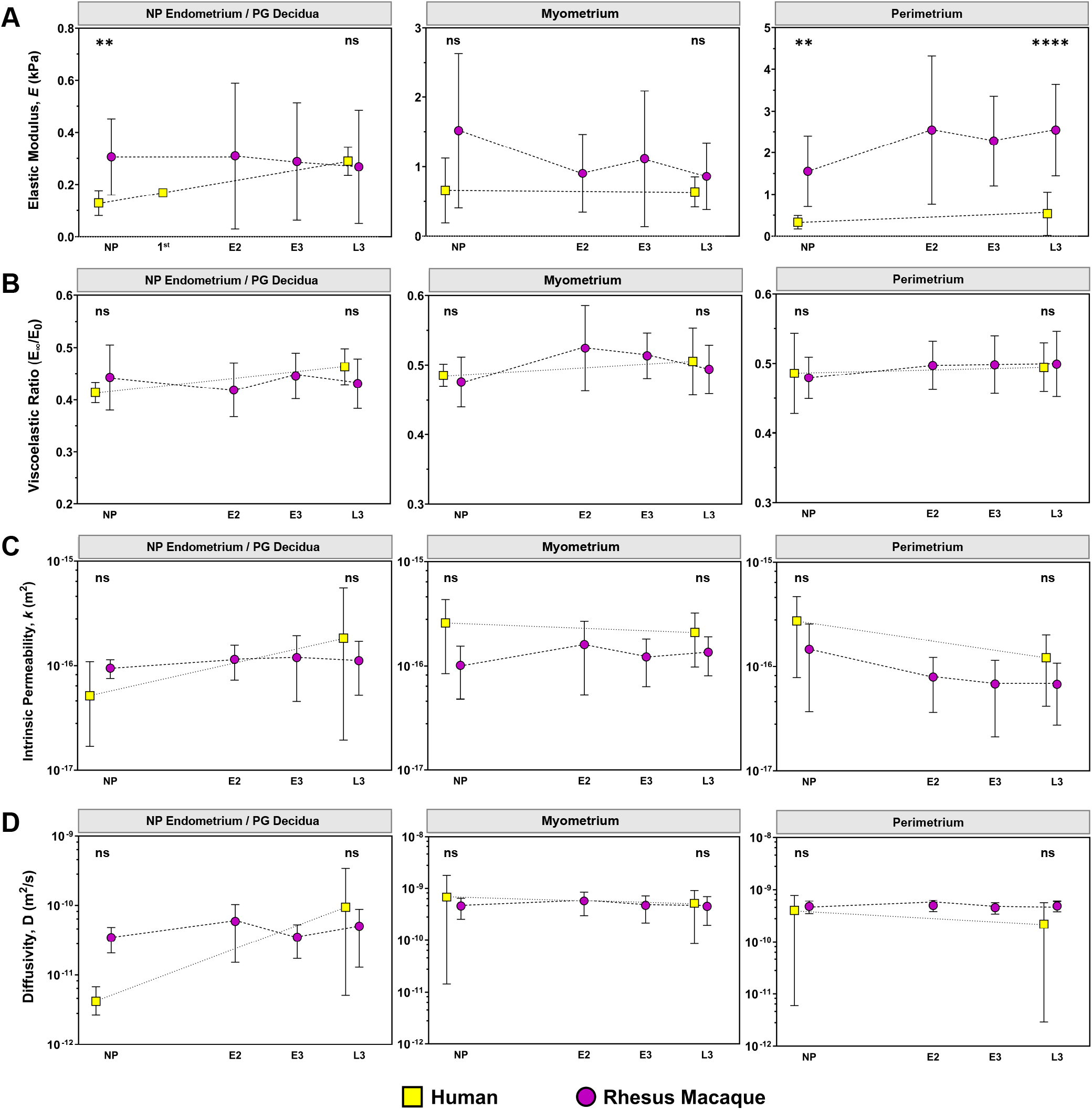
Comparison of Rhesus Macaque and Human Micromechanical Uterine Properties Across Gestation. Rhesus macaque data represents all animal subjects (*n* = 3−5) in this study sampled at three anatomic regions (anterior, fundus, posterior). Human data was taken from Fodera et al. (2024) for nonpregnant and late third-trimester (PG-CS) uterine tissue layers (*n* = 4−7) at the anterior region and Abbas et al. (2019) for the first-trimester decidua parietalis tissue (*n* = 5). Statistical significance between NHP and Human data was computed for NP and L3 groups and is marked above the appropriate comparison. Statistical differences found between human NP and L3 groups, as previously determined in Fodera et al. (2024), are denoted with a ♦ symbol to the right of each comparison.

The elastic moduli of the endometrium, decidua, and perimetrium tissue layers were found to be the most different between humans and NHPs; no species-related differences in myometrium stiffness were detected (Fig. 6). In particular, the NP endometrium was significantly less stiff in humans compared to NHPs and underwent stiffening in human pregnancy in a rather linear fashion (Fig. 6). Similarly, the perimetrium was consistently less stiff in humans across all gestational groups compared to NHPs (Fig. 6). Additionally, elastic modulus measurements in NHPs showed greater variability than in humans, as reflected by larger standard deviations (Fig. 6).

## Discussion

Here, we evaluated the structure-function relationship of the NHP uterus from nonpregnancy to late pregnancy, investigating differences across tissue layers and mid-gestational time points. Specifically, this microindentation dataset, together with histological and biochemical analysis, highlights drastic differences in structure, composition, and time-dependent material properties across the three uterine layers: NP endometrium / PG decidua, myometrium, and perimetrium. Interestingly, although pregnancy induced clear structural and compositional changes to the uterus, particularly for the decidua and myometrium layers, such differences were not reflected by alterations to uterine material properties on the microscale.

This study revealed distinct time-dependent material properties across the three structurally and functionally unique uterine layers. In pregnancy, a pronounced stiffness gradient was identified across the uterine wall, spanning from the decidua to the perimetrium (Fig. 3). In nonpregnancy, however, this stiffness gradient was found to only occur between the endometrium and myometrium, as the perimetrium was not mechanically distinct from that of the myometrium (Fig. 3). Mechanical gradients are functionally relevant in normal homeostasis and disease pathogenesis as they are able to modulate tissue growth and cell-cell signaling through mechanotransductive pathways^32–34^. Further, these mechanical gradients can influence fluid migration patterns and alter drug delivery efficacy through heterogeneous tissues^34,35^. Differences in the relationship between elastic modulus and intrinsic permeability were also found to occur in this dataset on a layer-specific basis (Fig. 5). We posit that the distinct clustering of the data points on the elastic modulus versus permeability plots is indicative of the heterogeneous composition of cellular and ECM components within the three tissue layers, and therefore, represents a unique, biophysical fingerprint for each. This connection is supported by work conducted by Islam and Oyen (2021) demonstrating a linear relationship between elastic modulus and permeability on a log-log scale for chemically cross-linked hydrogels^36^. Cross-link mobility, sample hydration, and polymer fraction have been shown to influence this relationship in hydrogels which scales with gel concentration (c)^36,37^: *k* ∝ *E*^−3*/*4^ ∝ *c*^−3*/*2^. Still, additional investigation is needed to precisely identify the cellular and ECM components contributing to these unique data clusters describing uterine mechanical properties.

This robust microindentation dataset further adds to fundamental understanding of pregnancy biomechanics by investigating how each of the three uterine layers evolves across mid-gestational time points in NHPs. Consistent with trends previously reported for human uterine tissue^20^, NHP pregnancy brought about no differences in the values of elastic modulus, viscoelastic ratio, permeability, or diffusivity for the myometrium and perimetrium at the microscale in small strain (Fig. 3, 4). The absence of material property changes across gestation in each uterine layer is thought to be a reflection of the small-strain regime evaluated in this study. The mechanical properties of nonpregnant and pregnant NHP uterine tissues were measured within the linear elastic regime (∼ 5% strain) of mechanical behavior at a length scale similar to that experienced by individual cells. This mechanical testing approach subjects samples to a complex loading profile of compression, radial tension, and shear^38^. Under compression, the mechanical response of biological tissues is largely dictated by the properties of their nonfibrillar ground substance, which is provided, in large part, by glycosaminoglycans (GAGs), proteoglycans, and fluid^39,40^. The fiber network alone cannot sustain compression but does constrain the lateral expansion of the ground matrix^39^. Engagement of the fiber network primarily occurs under tension through fiber uncrimping, alignment, and sliding^39–41^. Recent work has demonstrated that softening of the pregnant human myometrium is only observed under tensile loading at strains above 30%; no differences were observed under indentation up to 45% strain^20–22^. Therefore, this asymmetry in mechanical behavior under modalities of indentation and tension highlights fundamental differences in the contribution of the fiber network and ground substance to the overall material behavior of myometrium tissue^22,40^. Since only tensile testing at larger strains reveals softening of the human myometrium in late pregnancy, collagen fiber engagement appears to be necessary for observing this material behavior in pregnancy. Given the overlap in material properties for the human and NHP myometrium noted in this study, we posit that the NHP will exhibit similar tension-compression asymmetry and softening under tension in the large-strain regime. Aside from the myometrium, notable interspecies differences were identified for the nonpregnant endometrium and perimetrium. The baseline stiffness of the NP endometrium and perimetrium was greater for NHPs compared to humans (Fig. 6). Greater variability in the material properties of NHPs was also noted in this study (Fig. 6). In part, this increased mechanical heterogeneity can be attributed to differences in the number of anatomic regions sampled – three for NHPs and one for humans – but, still, intrinsic species differences cannot be excluded (Fig. S2). Notably, NHP pregnancy did not induce stiffening of the PG decidua relative to the NP endometrium, in contrast to the mechanical changes observed in human tissues (Fig. 3, 6). Such mechanical differences may be indicative of fundamental biological variations in the initiation and development of pregnancy in humans and NHPs at the maternal-fetal interface. Compared to humans, Rhesus macaques are known to exhibit superficial trophoblast invasion, undergo spiral artery remodeling at an earlier stage in pregnancy, and form a secondary placental disc^42^. The degree to which these processes individually shape the mechanics of the maternal decidua is presently unknown.

On a structural and compositional basis, notable differences were observed across all three uterine tissue layers both qualitatively and quantitatively (Fig. 2). At each gestational age evaluated, the organization of cellular and ECM components, as well as hydration measurements, varied across uterine layers; no significant changes in blood vessel area fraction were noted (Fig. 2). The effect of pregnancy on the structure and composition of all uterine layers was found to be rather mixed in this study. Notably, the L3 myometrium displays a shift in the relative proportion of smooth muscle and collagen components within a *mm*^2^ tissue area (Fig. 2). Similar increases in the proportion of smooth muscle to collagen content as a result of pregnancy have also been observed in humans^20^. In addition, a unique histological feature of focal edema was observed in the late third-trimester NHP uterus (Fig. 2). The increased interstitial spacing of the tissue’s microstructure is indicative of increased swelling of the myometrium in the late third trimester of pregnancy. Such changes in fluid homeostasis may be a result of shifts in microvascular pressure^43^, electrolyte concentrations^44^, and composition of hydrophilic ECM proteins (e.g., proteoglycans and hyaluronan)^45,46^. Interestingly, this phenomenon is not reflected by quantitative tissue hydration measurements reported in this study which show no change in the overall hydration of the myometrium with pregnancy (Fig. 2). This disparity is likely due to length-scale effects, as the *µm*^2^ edema feature represents a small portion of the overall *mm*^2^ tissue sample measured. Experimental limitations such as flash-freezing could produce histological artifacts, but the lack of similar features observed in NP, E2, and E3 tissues subjected to the same processing conditions weakens this argument. On the matter of vascular remodeling, no pregnancy-induced changes to blood vessel area fraction were observed for both the decidua and myometrium (Fig. 2). The absence of notable vascular differences, particularly in the decidua, is likely due to sampling from the parietalis region, as the decidua basalis undergoes more pronounced vascular remodeling during pregnancy^47^. Still, studying vascular remodeling in this region is valuable as increased blood vessel density in first-trimester human decidua parietalis tissue has been associated with spontaneous abortions^48^. It is important to note that the analysis of compositional and structural features in this study does not encompass all cellular and ECM components (e.g., elastin, proteoglycans) that may influence the mechanical behavior of the uterus and such structures warrant further investigation.

Taken together, these results highlight the validity of Rhesus macaques as a model for studying certain aspects of human uterine biomechanics and mechanobiology, both to advance foundational knowledge of pregnancy and to guide therapeutic innovation for obstetric conditions. The myometrium exhibited the most notable interspecies similarities in terms of mechanics and structure (Fig. 2, 3, 6). Since the myometrium layer is largely responsible for the growth experienced by the uterus, deleterious alterations to the mechanical and structural properties of this tissue can impede normal uterine stretch and vascular remodeling. Therefore, the ability to study uterine growth and remodeling *in vivo* at time-points that are ethically restricted in human pregnancy is critical for elucidating the pathophysiological features of obstetric conditions, including miscarriage, preterm birth, preeclampsia, uterine rupture, and fetal growth restriction. While the myometrium exhibited broad interspecies similarities, notable differences were identified in the baseline mechanical properties of the nonpregnant endometrium and perimetrium, as well as in their pregnancy-associated changes. Therefore, this highlights limitations in the use of Rhesus macaques as an appropriate model to study mechanically driven interactions at the maternal-fetal interface. In fact, all decidua tissues analyzed in this study represent the parietalis region (i.e., distant from placentation) and not the basalis region (i.e., at the site of placentation), so the findings may not reflect certain region-specific adaptations critical to placentation, such as vascular remodeling. Further supporting this point, Abbas et al (2019) documented mechanical differences between decidua parietalis and decidua basalis tissues in first-trimester human pregnancies, which were attributed, in part, to differences in spiral artery remodeling^19^. Nonetheless, this study reinforces the utility of the Rhesus macaque as a valid model for investigating certain aspects of uterine biomechanics and mechanobiology in pregnancy.

While this study provides important insights into mechanical and structural changes of the NHP uterus, several methodological limitations should be considered when interpreting the findings. First, uterine tissues underwent flash-freezing at the time of tissue collection and underwent one to two freeze-thaw cycles prior to analysis. It is well understood that flash freezing often causes ice crystal formation in biological samples, damaging cell membranes and leading to poor cell viability^49^. However, the ECM, which is largely responsible for dictating the mechanical properties of biological tissues, undergoes minimal changes to its structure as a result of freezing^50,51^. In the literature, the degree to which freeze-thaw cycles alter the mechanical properties of biological tissues is variable^22,52–56^. Some studies have demonstrated the capacity of multiple freeze-thaw cycles to alter mechanical properties^52–54^, while other studies observed no effects^22,55,56^. Whether mechanical changes occur due to freeze-thaw cycles appears to be largely dependent on the specific tissue in question and the mechanical method of characterization. Most notably, recent work from our group published in Fang et al (2025) demonstrated no change in the mechanical response for late third-trimester pregnant human uterus tissue subjected to macro-indentation^22^. Therefore, future studies are needed to elucidate the exact effect of freeze-thaw cycles on nonhuman primate uterine layers in nonpregnant and pregnant states. Second, a key limitation of this study is the small number of animal subjects (*n* = 3 − 5 / gestational group) represented, reducing the statistical power and, potentially, the generalizability of our findings. We were largely constrained by the availability of Rhesus macaques from the Wisconsin National Primate Research Center at the time this study was conducted during the peak of the COVID-19 pandemic. To address this limitation in our mechanical data, tissues were sampled at three anatomic regions, and this data was pooled on an individual animal basis for statistical analysis. However, the extent of regional variations in tissue composition and structure throughout the pregnant uterus remains unknown. Further, the degree to which confounding variables such as age, menstrual cycle stage, and number of previous pregnancies (i.e., gravidity) alter the mechanical and structural properties of the uterus is unknown and was not accounted for in this study. Future work is needed to tease out the effect of these patient-specific variables. Lastly, while microindentation is a powerful tool to measure the time-dependent material properties of biological tissues spatially, the physical limits of this technique must be considered. Microindentation spatially maps a *mm*^2^ region of tissue, which is considerably smaller than the entire *cm*^2^ surface area of intact uterine tissue *in vivo*. In particular, the permeability measurements reported in this study do not capture the larger interconnected pore network likely present in these tissues^57^. While no studies to date have directly measured the permeability of the uterus *a priori*, the hydraulic permeability of human cervix tissue measured with a passive pressure gradient^58^ exhibited values several orders of magnitude greater than that which is reported for the NHP and human uterus with microindentation^20^.

Despite the aforementioned limitations, this study establishes the normal heterogeneity of material, structural, and compositional properties across the NHP uterine layers in nonpregnant and pregnant states, revealing notable similarities to the human uterus, particularly for the myometrium tissue layer. Characterizing baseline changes that occur throughout healthy pregnancies in a physiologically comparable NHP animal model is foundational to better understanding and predicting healthy and disordered alterations in human gestation with *in vitro, in vivo*, and *in silico* approaches.

## Methods

### Animal Study and Tissue Collection

The study details reported in this manuscript abide by the ARRIVE guidelines^59^. This protocol was approved by the University of Wisconsin-Madison Institutional Animal Care and Use Committee (IACUC) and adhered to federal guidelines and regulations for proper animal care and experimentation. Nonpregnant (NP) and pregnant (PG) Rhesus macaques of the female sex were recruited from the Wisconsin National Primate Research Center (WNPRC) and chosen from among those scheduled for removal from the breeding colony within 1–2 years. Specifically, this study investigated Rhesus macaques in nonpregnancy (n = 3) and at three time points in pregnancy: early 2^nd^ (E2, n = 3), early 3^rd^ (E3, n = 3), and late 3^rd^ (L3, n = 5) trimesters (Fig. 1). All animal subjects overlap with those reported in Fang et al (2024)^60^. Detailed subject information, including age, gestational age, weight, body condition score, gravidity, placenta location, and past obstetric history, are noted in Table 1. To note, body condition score (BCS) is a semi-quantitative assessment of an animal’s lean body mass evaluated on a 1.0 to 5.0 scale, where 3.0 represents the optimal body condition score^61^. Weight was recorded at the time of surgery. Gravidity represents the total number of previous pregnancies and does not include the current pregnancy for PG subjects.

**Table 1.** Detailed summary of individual non-human primate characteristics. VD ≡ Vaginal delivery, CS ≡ Cesarean section.

| Animal ID | Age (yrs) | Gestational Length (days) | Gravidity | Animal Weight (kg) | Body Condition Score | Placenta Disc Locations (Primary & Secondary) | Prior Obstetric History |
| --- | --- | --- | --- | --- | --- | --- | --- |
| NP-1 | 4.1 | N/A | 0 | 5.8 | 3 | N/A | None |
| NP-2 | 15.2 | N/A | 5 | 13.41 | 4.5 | N/A | VDx4, CSx1 |
| NP-3 | 11.8 | N/A | 5 | 8.14 | 4 | N/A | VDx4 |
| E2-1 | 15.2 | 72 | 7 | 9.07 | 3.5 | Unknown | VDx4, CSx3 |
| E2-2 | 8.9 | 71 | 3 | 8.44 | 3 | Posterior Lateral & Anterior Lateral | VDx2, CSx1 |
| E2-3 | 11 | 73 | 1 | 6.01 | 3 | Anterior & Posterior | VDx1 |
| E3-1 | 8 | 110 | 3 | 10.96 | 3.5 | Anterior & Posterior | VDx3 |
| E3-2 | 18 | 112 | 8 | 9.48 | 3 | Fundo-Anterior & Posterior | VDx5, CSx3 |
| E3-3 | 18.8 | 119 | 4 | 12.6 | 4 | Anterior & Posterior | VDx1, CSx3 |
| L3-1 | 16 | 161 | 8 | 11.12 | 3.5 | Anterior Lateral & Posterior Lateral | VDx5, CSx3 |
| L3-2 | 9.7 | 158 | 4 | 10.83 | 3.5 | Anterior & Posterior-Fundus | CSx4 |
| L3-3 | 18.4 | 152 | 4 | 11.77 | 4.5 | Anterior Lateral & Posterior Lateral | VDx1, CSx1 |
| L3-4 | 12 | 157 | 5 | 6.11 | 3 | Anterior & Posterior | VDx3, CSx2 |
| L3-5 | 14.2 | 154 | 4 | 8.54 | 3.5 | Unknown | VDx2, CSx2 |

All subjects underwent a total hysterectomy procedure to remove the uterus and cervix, which was performed by a board-certified obstetrician-gynecologist (H.F.) and overseen by WNPRC veterinarians and staff. This was a terminal study, and all dams and pre-viable fetuses were euthanized in accordance with humane standards set forth by the American Veterinary Medical Association (AVMA). Hysterectomies were performed on NP, E2, and E3 subjects following euthanasia and on L3 Rhesus macaques prior to euthanasia to allow for fetal survival. NP, E2, and E3 animals received ketamine hydrochloride (15 mg/kg, intramuscular [IM]) for sedation and sodium pentobarbital (50 mg/kg, intravenous [IV]) for euthanasia; no analgesics were given. Prior to surgery, L3 animals were sedated with ketamine hydrochloride (15 mg/kg, IM) and propofol (1-5 mg/kg, IV) and received lidocaine (2 mg/kg) and buprenorphine hydrochloride (0.05 mg/kg) epidurally for analgesia. Food was only restricted if recommended by the veterinarian prior to anesthesia, with the maximum fast being 24 hours. Once sedated, L3 subjects were intubated with a cuffed endotracheal tube and the surgical site was clipped of fur and sterilely prepped. General anesthesia was maintained for L3 subjects during the course of the hysterectomy procedure with a constant rate infusion of propofol (0.1-0.6 mg/kg/min, IV). Adequate anesthesia level was assured by checking reflexes and monitoring for any spontaneous movement. During surgery, heart rate, respiration rate, and blood oxygen were monitored every 5-10 minutes for L3 subjects, and animals received routine isotonic intravenous fluids (5-10 ml/kg/hr) to avoid dehydration. Sterile instruments were used for all procedures. A midline abdominal skin incision was made from 4 cm cranial to the umbilicus to just cranial to the pubis. The caudal abdominal wall was incised to expose the uterus. The uterus was exteriorized and surrounded with sterile, saline-soaked lap sponges. For PG subjects, a sharp incision (5-6 cm) was made through all layers of the uterus at the fundus region and the amniotic sac, and the fetus was subsequently removed through the cesarean incision. All L3 fetuses were immediately transferred to a neonatal incubator and cared for by WNPRC staff. The fundal incision on the uterus was sutured closed with 3-0 Vicryl. Once hemostasis was assured, the maternal uterus and cervix were then transected from the abdominal cavity and placed on a tray for subsequent dissection by a member of the research team (S.F.). The abdominal incision was sutured closed with 3-0 Vicryl using a simple continuous pattern. L3 subjects were subsequently euthanized with an overdose of sodium pentobarbital (50 mg/kg, IV). The overall duration of this procedure and anesthesia use was no more than 120 minutes.

Full-thickness uterine wall segments were dissected from three anatomic regions (i.e., anterior, fundus, and posterior) and contained all three tissue layers (i.e., endometrium/decidua, myometrium, and perimetrium). Samples were flash-frozen on dry ice and stored at −80ºC. Uterine samples were then shipped on dry ice from the University of Wisconsin-Madison to Columbia University per a material transfer agreement and once again stored at −80ºC until further processing.

### Histology

For each NHP subject, uterine cross-sections, which contained all three tissue layers, were prepared for histology; only one anatomic region per subject was included, as noted in Table S1. Samples were fixed in 10% formalin solution for 24 hrs and subsequently transferred to 70% ethanol solution. Samples were paraffin-embedded and sectioned to a thickness of 5 µm by the Molecular Pathology Core Facilities at Columbia University Irving Medical Center (CUIMC). To observe histomorphology, all samples were stained for Hemotoxylin & Eosin (H&E) and Masson’s Trichrome using standard protocols^62^. Samples were imaged under brightfield microscopy with a Leica Aperio AT2 whole slide scanner up to 20x magnification and visualized with the Aperio ImageScope software (v12.3.1.6002, Leica Microsystems, Wetzler, Germany). Scanned histology slides have been made publicly available on Columbia University’s Academic Commons^63^. All slides were reviewed by a board-certified pathologist (X.C.) who specializes in gynecologic pathology and cytopathology. Histological assessment of uterine pathology and estimated menstrual stage is noted in Table S1.

#### Image Quantification

The relative proportions of collagen and smooth muscle content in the myometrium were quantified from Masson’s Trichrome stained tissue. Three representative images per NHP subject were taken at 10x magnification with a Leica DMi1 Inverted Microscope using the Leica Application Suite X (LAS-X). For this quantification, regions containing blood vessels in more than fifty percent of the image area were avoided. The areas of blue and red color, corresponding to collagen and smooth muscle content, respectively, were quantified in ImageJ (NIH, Bethesda, MD, USA) with RGB color deconvolution and a thresholding function.

Additional analysis was conducted to quantify the number and size of blood vessels in the endometrium, decidua, and myometrium tissue layers for each animal subject. Blood vessels were manually identified on Masson’s Trichrome stained uterine cross sections using Aperio ImageScope’s annotation tool. The stained portion of each tissue layer was analyzed in its entirety. The collective areas (*µm*^2^) of blood vessels were computed and divided by the total tissue layer area to determine the percent blood vessel area.

### Tissue Hydration

Lyophilization was used to determine tissue hydration of all NHP uterine layers using a FreeZone 4.5 Liter Benchtop Freeze Dry System (Labconco, Kansas City, MO). For each tissue layer, three tissue samples per animal subject were analyzed. All samples were taken from the posterior region of the uterus and dissected into small (*mm*^3^) pieces. Wet and dry sample weights were measured in a pre-weighed 1.5 ml Eppendorf tube before and after lyophilization using an analytical balance (MS105, Mettler Toledo, Greifensee, Switzerland) with 0.01 mg readability. Tissue hydration was calculated with the following equation:

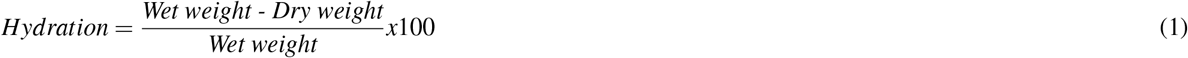

### Microindentation Testing

Spherical microindentation (Piuma, Optics11Life, Amsterdam, NE) was utilized to determine the material properties of uterine tissue. A 50 µm probe radius with a cantilever stiffness of 0.15 – 0.5 N/m was used. In preparation for testing, samples were dissected, adhered to a glass dish with superglue (Krazy Glue, Atlanta, GA), and thawed at 4°C overnight in 1X PBS solution supplemented with 2 mM ethylenediaminetetraacetic acid (EDTA). This swelling protocol was based on previous experiments conducted on human uterine tissue by Fang et al. (2022)^21^. Immediately prior to testing, the sample was equilibrated to room temperature for 30 minutes and subsequently tested in Opti-free contact lens solution (Alcon, Fort Worth, TX, USA) to reduce adhesion between the glass probe and sample^64^. Tissues were indented to a fixed depth of 4 µm under indentation control, corresponding to a 5% indentation strain and contact area of 380 µ*m*^2^. Following a 2 s ramp to the prescribed indentation depth, the probe’s position was held for 15 s to yield a load relaxation curve approaching equilibrium. All tissues were at least 1 mm thick and tested within two freeze-thaw cycles.

#### Individual Uterine Tissue Layers

For each gestational group (i.e., NP, E2, E3, L3), the material properties of all three individual uterine tissue layers (i.e., endometrium/decidua, myometrium, and perimetrium) were assessed with microindentation. For each animal subject, each uterine layer was measured at three different anatomic regions (i.e., anterior, fundus, and posterior). To ensure reliable measurements taken for thinner tissue layers, the surface of the endometrium-decidua, and perimetrium were directly tested. The orientation of the myometrium was variable and not explicitly noted. Given that the size and geometry of the tissues were so irregular, the number of indentation points also varied; approximately 100 points were measured per sample to capture sufficient intra-sample variability. The distance between individual indentations was kept constant at 200 µm.

#### Uterine Wall Cross Sections

Spatial stiffness variations were assessed across the entire length of the posterior uterine wall tissue section from the perimetrium to the endometrium-decidua. One subject per gestational group was assessed. The width of the tested region was kept constant at 1mm and the distance between individual indentations was fixed at 200 µm.

### Microindentation Data Analysis

#### Poroelastic-Viscoelastic (PVE) Model

For individual tissue layers, load versus time data from the hold portion of the indentation protocol were fit with a combined poroelastic-viscoelastic (PVE) model in Matlab based on an established analytical solution with a nonlinear least-squares solver^36,65,66^. The coupled effect of the material’s poroelastic (*P*_*PE*_) and viscoelastic (*P*_*VE*_) force responses is described by:

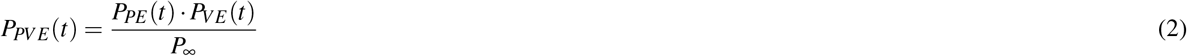

The viscoelastic force response is calculated using a generalized Maxwell model, consisting of a linear spring connected in parallel with two Maxwell units, each containing a linear spring and dashpot connected in series. The viscoelastic component of the model is defined by the following equation:

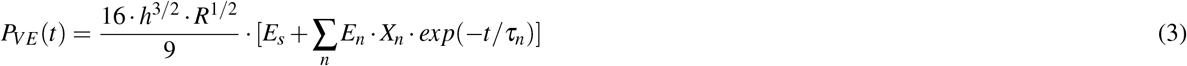

where *E*_*s*_ and *E*_*n*_ are the elastic moduli of the linear spring and the *n*^*th*^ Maxwell element (n = 2), respectively, *h* is the applied indentation depth, *R* is the probe radius, and *τ*_*n*_ is the characteristic relaxation time of the *n*^*th*^ Maxwell element (n = 2). A ramp correction factor (*X*_*n*_ = (*τ*_*n*_*/t*_*r*_) · [*exp*(−*t*_*r*_*/τ*_*n*_) − 1]) is included to account for the two-second ramp time (*t*_*r*_) since the original Maxwell model assumes a step loading function^67^. Instantaneous elastic modulus (*E*_0_) and equilibrium elastic modulus (*E*_∞_) parameters are determined from Eqn. 3 when *t* = 0 and *t* = ∞, respectively.

The poroelastic force response is calculated from the analytical solution published in Hu et al. 2010:

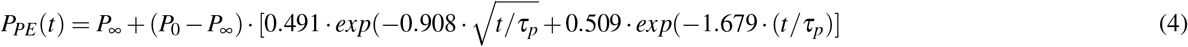

*P*_0_ is the initial force at the beginning of the load relaxation curve and is calculated from the model defined in Hu et al. 2010 as *P*_0_ = (16*/*3)· *G*_*PE*_ · *R*^1*/*2^ · *δ*_0_^3*/*2^, where *G*_*PE*_ is the apparent poroelastic shear modulus. *P*_∞_ is the estimated equilibrium force given by *P*_∞_ = *P*_0_*/*[2(1 − *ν*_*d*_)], where *ν*_*d*_ is the drained Poisson’s ratio. *τ*_*p*_ is the poroelastic time constant in the defined as *τ*_*p*_ = *a*^2^*/D*, where *a* is the indentation contact radius 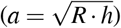 and *D* is diffusivity. Intrinsic permeability (*k*) is calculated as follows:

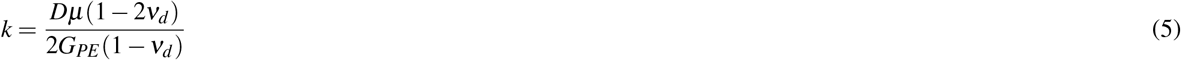

The interstitial fluid viscosity (*µ*) is assumed to be equivalent to the dynamic viscosity of water at 25°C (*µ* = 0.89 × 10^−3^*Pa* · *s*). Material incompressibility is assumed, wherein material volume does not change under applied deformation, and therefore, undrained Poisson’s ratio (*ν*) is set as 0.5. The apparent poroelastic modulus (*E*_*PE*_) is calculated from apparent poroelastic shear modulus as *E*_*PE*_ = 3*G*_*PE*_.

Fitted data points were excluded from the final data set if the load relaxation curve displayed (i) sharp discontinuities, (ii) increasing loads over time, or (iii) ΔP ∼ (*P*_*max*_ – *P*_*min*_) less than 0.005 *µ*N.

#### Hertzian Contact Model

Data for uterine wall cross-sections were analyzed with the Hertzian contact model to reduce the number of points removed due to exclusion criteria. The apparent elastic modulus (*E*_*r*_) was determined by fitting each load versus indentation curve from the initial loading portion of the indentation protocol with the following equation^31,68^:

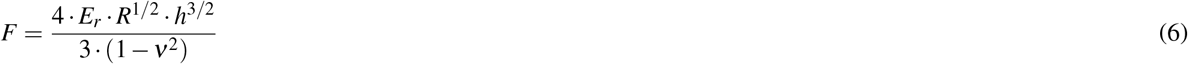

where *F* is the applied force, *R* is the probe radius, *h* is the applied indentation depth, and *ν* is Poisson’s ratio. A value of 0.5 is prescribed for *ν* to align with incompressibility assumptions. This model assumes contact between a sphere and a half-space for a material that is linear elastic. Data fitting was performed with a customized code in Matlab (Mathworks, Natick, MA, USA) using a nonlinear least-squares solver, identical to what has been previously published in the literature^20,36^. Data points were excluded if the corresponding *R*^2^ value was less than 0.5, indicating a poor model fit.

### Statistical Analysis

Statistical analysis was performed using RStudio (v1.3.1056) or GraphPad Prism (v.10.0.2). Normality of all data was first assessed with Q-Q plots. In instances of non-normal data distributions, data were normalized with a logarithmic transformation. A linear mixed-effects model was employed to analyze all datasets in this study which investigated differences across tissue layers or gestational groups. In all cases, animal ID was set as the random variable. Multiple comparisons were assessed with a Tukey post-hoc test. To assess differences in material properties between NHPs and humans^19,20^, an unpaired t-test with a Welch correction was performed for each parameter at NP and L3 time points. To determine the inter-correlation of material properties by tissue layer, Pearson correlation coefficients were calculated. Linear regression analysis was performed in noted cases of continuous variables; the multiple *R*^2^ and *p* values are reported for each fit. Significance was set at a 95% confidence level for all analyses. P-value symbols are defined as follows: ^*^ *p<*0.05, ^**^ *p<*0.01, ^***^ *p<*0.001, ^****^ *p<*0.0001. A .xlsx file summarizing the p-values from all statistical comparisons has been included in supplementary material.

## Supporting information

P-values of Statistical Tests

## Acknowledgements

The research was supported in part by the National Science Foundation (NSF) Graduate Research Fellowship (DGE-2036197) to DMF, The Iris Fund, the Eunice Kennedy Shriver National Institute of Child Health Human Development Grant R01HD072077 to TH, IRM, HF, and KM, and the Office Of The Director, NIH Award P51OD011106 to the Wisconsin National Primate Research Center, University of Wisconsin-Madison. We would also like to thank Qi Yan of Columbia University Irving Medical Center for statistics consultation and Michele Schotzko of the Wisconsin National Primate Research Center for confirmation of animal study details. Figures for this paper were created with BioRender.com, Adobe Illustrator, and GraphPad Prism under registered academic licenses.

## Author Contributions Statement

Conceptualization: DMF, KMM

Methodology: DMF, KMM, MLO, XC

Investigation: DMF, EZX, MW

Formal Data Analysis: DMF, EZX

Data Interpretation: DMF, EZX, CADC, MLO, XC, KMM

Resources: SF, TH, IRM, HF, KMM, JYV

Writing, Original Draft: DMF

Writing, Review and Editing: DMF, EZX, CADC, XC, MLO, IRM, JYV, TH, HF, KMM

Visualization: DMF

Supervision: KMM

Funding Acquisition: TH, IRM, HF, KMM

## Competing Interests Statement

The authors declare that they have no competing interests.

## Data and materials availability

All quantitative data (https://doi.org/10.7916/bb8j-pm63) and scanned histological images (https://doi.org/10.7916/w6nr-5x38) have been made available on Columbia University’s Academic Commons. Codes used for all data analyses are available upon request to the corresponding author (KMM).

## Notes

https://academiccommons.columbia.edu/doi/10.7916/w6nr-5x38

https://academiccommons.columbia.edu/doi/10.7916/bb8j-pm63

### Competing Interest Statement

The authors have declared no competing interest.

### Summary of Updates

The discussion section has been revised. Additional figures have been added to the supplementary materials.

