## Supplementary Figures & Tables for "Time-Dependent Material Properties and Composition of the Nonhuman Primate Uterine Layers Through Gestation"

### Supplemental Figures & Tables

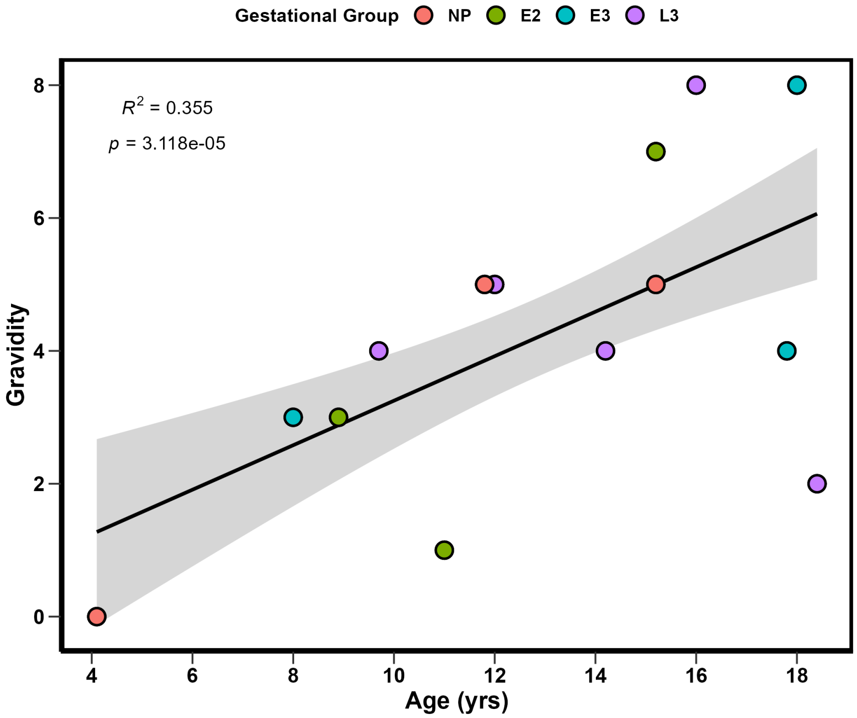


**Figure S1.** Correlation between age and gravidity for animal subjects studied in this cohort. $R^2$ and $p$ values are noted.

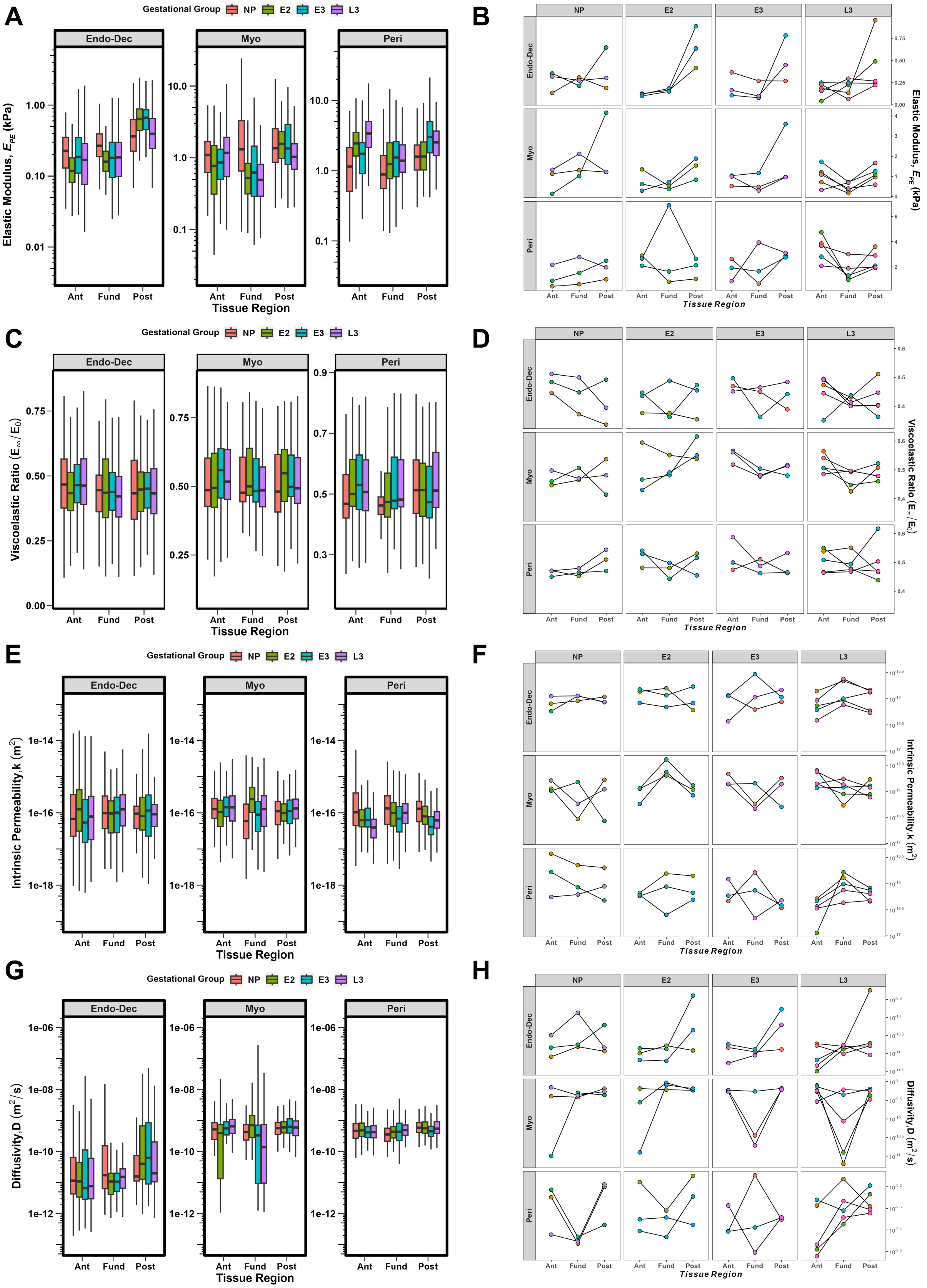


**Figure S2. Variations in Material Properties Across Anatomic Regions (Anterior, Fundus, Posterior)**. Elastic modulus (A,B), viscoelastic ratio (C,D), intrinsic permeability (E,F), and diffusivity (G,H). All data for a given tissue layer, gestational group and anatomic region are shown as box and whisker plots (left column). The right column depicts matched values for each tissue layer and gestational group for a given animal across the three anatomic regions. Each point represents the median value of all indentation points measured for a single sample.

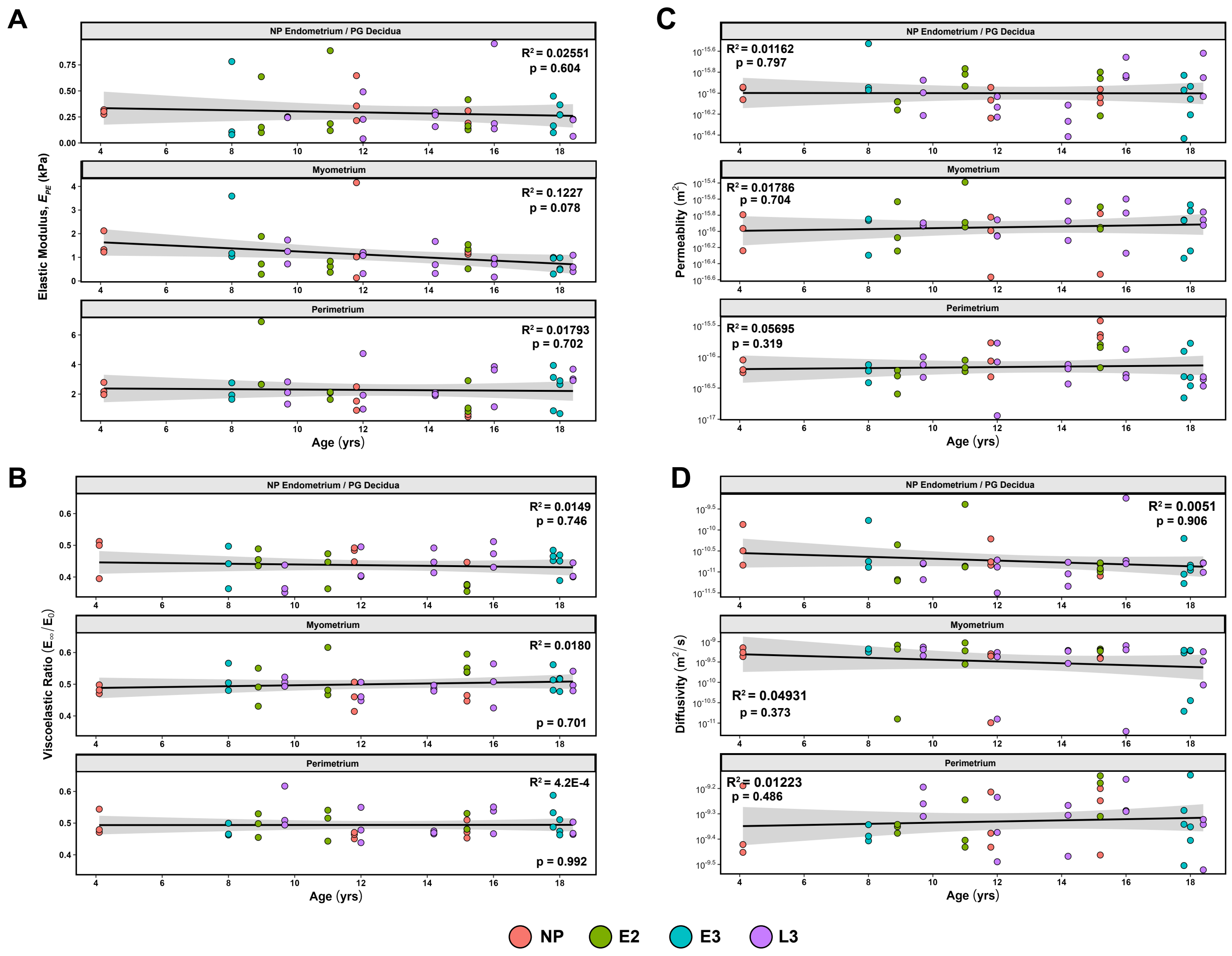


**Figure S3. Correlation of Uterine Layer Mechanical Parameters with Animal Age.** Each point represents the median value of all indentation points measured for a single sample. All animal subjects ($n = 3 - 5$) in this study sampled at three anatomic regions (anterior, fundus, posterior) are represented in these data. The $R^2$ and $p$ values for each correlation are noted. Standard deviations are indicated by the shaded grey areas.

| Animal ID | Anatomic Region Analyzed | Estimated Menstrual Cycle Stage | Additional Pathological Findings |
|---|---|---|---|
| NP-1 | Anterior | Proliferative | Nodular and disorganized myometrium |
| NP-2 | Anterior | Unknown – Basalis tissue only | Fibrous basalis tissue; Fibrosis of serosa |
| NP-3 | Anterior | Late Proliferative / Early Secretory | Adenomyosis |
| E2-1 | Anterior | N/A. Decidua Parietalis | None |
| E2-2 | Anterior | N/A. Decidua Parietalis | None |
| E2-3 | Anterior | N/A. Decidua Parietalis | None |
| E3-1 | Anterior | N/A. Decidua Parietalis | None |
| E3-2 | Fundus | N/A. Decidua Parietalis | Endosalpingiosis; Notable fibrotic scar tissue that extends vertically from serosa to decidua – indicative of a previous C-section incision |
| E3-3 | Anterior | N/A. Decidua Parietalis | Focal thickening of serosa |
| L3-1 | Anterior | N/A. Decidua Parietalis | None |
| L3-2 | Anterior | N/A. Decidua Parietalis | None |
| L3-3 | Anterior | N/A. Decidua Parietalis | None |
| L3-4 | Anterior | N/A. Decidua Parietalis | None |
| L3-5 | Anterior | N/A. Decidua Parietalis | None |

**Table S1.** Summary of histological findings for all NHP subjects with estimated menstrual cycle stage for NP individuals.

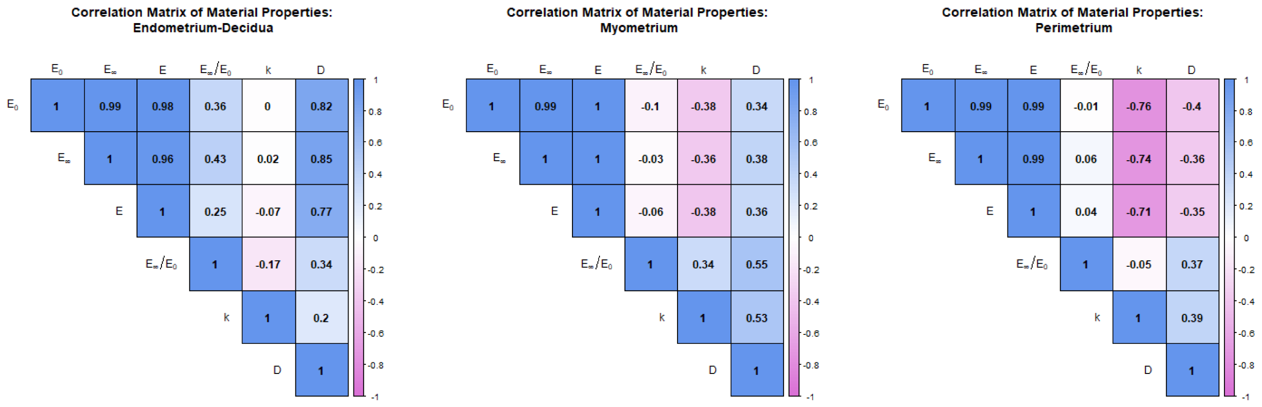


**Figure S4. Inter-correlation of Material Parameters by Uterine Tissue Layer.** Values close to -1, shown in pink, indicate a strong negative correlation between the two parameters, while values close to 1, shown in blue, indicate a strong positive correlation.

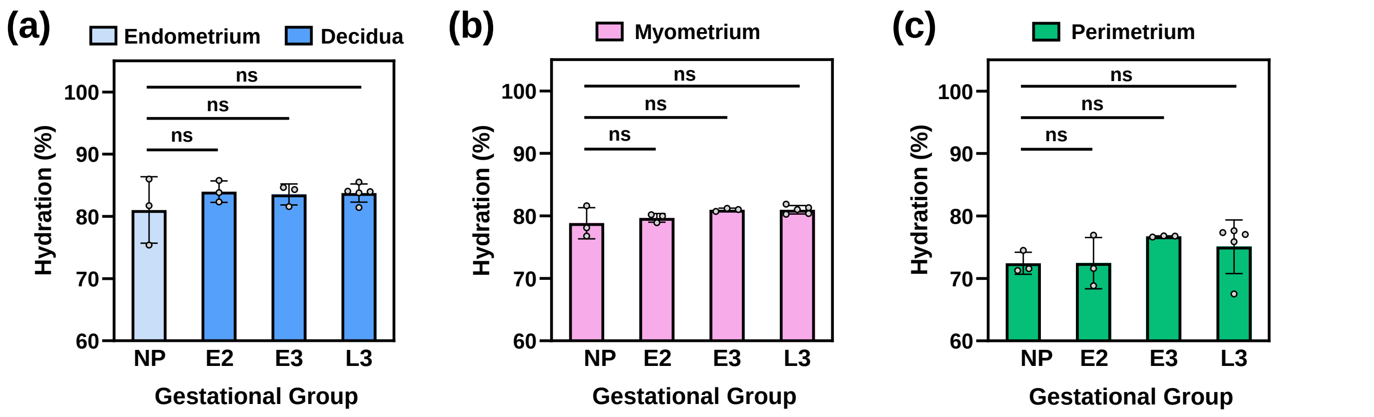


**Figure S5.** **Hydration (%) of individual NHP uterine layers throughout pregnancy.** (a) NP endometrium and PG decidua, (b) myometrium, and (c) perimetrium tissue layers sampled at the posterior region. Each symbol represents the average of three biological replicates from a single animal subject. All animal subjects (n = 3-5) in this study are represented in these data.

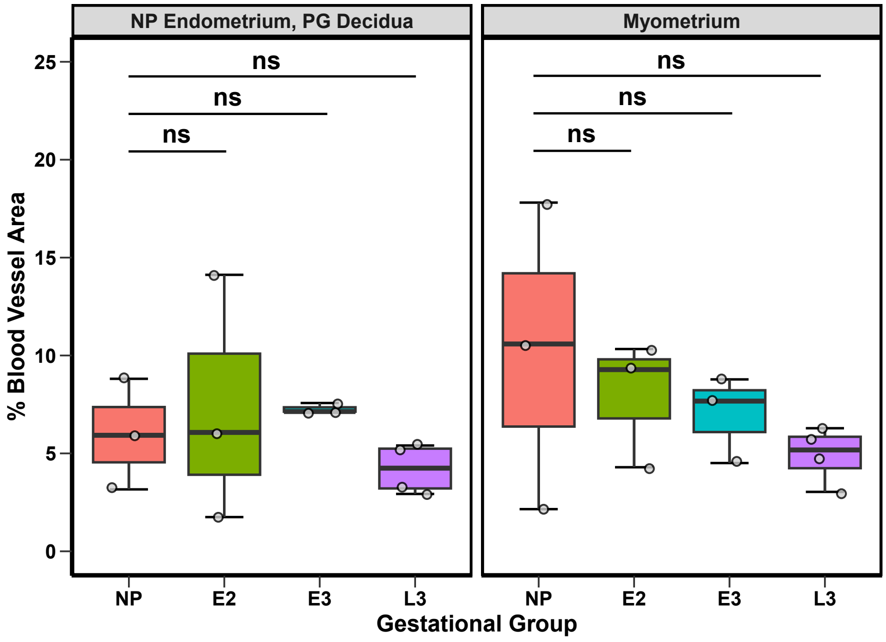


**Figure S6.** **Proportion of blood vessel area across gestation by tissue type.** No quantification was conducted for the perimetrium layer. All animals, except one L3 subject, are represented in these data.
